# Nell2 regulates the contralateral-versus-ipsilateral visual projection as a layer-specific positional cue

**DOI:** 10.1101/392597

**Authors:** Chizu Nakamoto, Elaine Durward, Masato Horie, Masaru Nakamoto

## Abstract

**SUMMARY STATEMENT:** Nell2 is an ipsilateral layer-specific axon guidance cue in the visual thalamus and contributes to establishment of the eye-specific retinogeniculate projection by specifically inhibiting contralateral retinal axons.

**ABSTRACT:** In mammals with binocular vision, retinal ganglion cell (RGC) axons from each eye project to eye-specific layers in the contralateral and ipsilateral dorsal lateral geniculate nucleus (dLGN). Although layer-specific axon guidance cues that discriminate contralateral and ipsilateral RGC axons have long been postulated as a key mechanism for development of the eye-specific retinogeniculate projection, the molecular nature of such cues has remained elusive. Here we show that the extracellular glycoprotein Nell2 (also known as Nel) is expressed in the dorsomedial region of the dLGN, which corresponds to the layer receiving ipsilateral RGC axons. In Nell2 mutant mice, contralateral RGC axons invaded the ipsilateral layer of the dLGN, and ipsilateral axons terminated in partially fragmented patches, forming a mosaic pattern of contralateral and ipsilateral axon termination zones. *In vitro,* Nell2 exerted inhibitory effects on contralateral, but not ipsilateral, RGC axons. These results provide evidence that Nell2 acts as a layer-specific positional label in the dLGN that discriminates contralateral and ipsilateral RGC axons, and that it plays essential roles in establishment of the eye-specific projection patterns in the retinogeniculate system.

## INTRODUCTION

In his treatise on optics, Isaac Newton predicted that binocular vision would rely on the convergence of information from both eyes on the same site in the brain, and that a partial decussation of the optic nerve fibres would be necessary for binocular integration (Newton, 1730). His anatomical prediction was verified not long afterwards in dissections of the mammalian visual system (Pettigrew, 1986; Polyak, 1957). In mammals with good binocular vision, visual information from each eye is transferred to both sides of the brain via axons of retinal ganglion cells (RGCs): RGC axons from the nasal retina cross the midline at the optic chiasm and project to the contralateral side of the brain, whereas axons from a segment of the temporal retina project to the ipsilateral side. This partial decussation of RGC axons allows integration of visual information from both eyes in the brain, thus underpinning disparity-based stereopsis (depth perception and visual measurement of distance) (Wilks et al., 2013). Ipsilaterally-projecting RGCs localise in the defined retinal area of binocular overlap, and their number and distribution correlate with the extent of binocular vision and the orientation of the orbits (Heesy, 2004). In mice, ipsilateral RGC axons arise from a ventrotemporal segment of the retina and comprise c.3-5% of the optic nerve fibres (Erskine and Herrera, 2014; Petros et al., 2008). In contrast, mammals without binocular overlap in their visual fields and all the non-mammalian vertebrates have a total decussation of the nerve fibres at the optic chiasm and thus no ipsilateral RGC fibres (O’Leary et al., 1983; Polyak, 1957).

In the mammalian visual system, RGC axons project in an orderly manner to their main forebrain target, the dorsal lateral geniculate nucleus (dLGN) of the thalamus. The patterning of retinogeniculate axons involves (1) segregation of right and left eye inputs, (2) topographic map formation, and (3) placement of eye-specific layers that locate in stereotypic positions in the dLGN (Huberman et al., 2008; Pfeiffenberger et al., 2005). In the strict sense, the term “layer” refers to a thin sheet of discrete cellular groupings. However, “eye-specific LGN layers” is also used to refer to the regions of stereotyped size, shape and position formed by RGC axons arising from each eye (Huberman et al., 2005). In this article, we use the term “layer” in this latter sense.

Spontaneous retinal activity has been thought to play crucial roles in eye-specific segregation of RGC axons in the dLGN (Huberman, 2007), and several molecules have been shown to play important roles in this activity-dependent segregation (Huh et al., 2000), including class I MHC (major histocompatibility complex), CD3ζ (Huh et al., 2000), neuronal pentraxins (Bjartmar et al., 2006), Zic2, and serotonin transporter (Sert) (Garcia-Frigola and Herrera, 2010).

Activity-dependent mechanisms, however, cannot account for the stereotypical locations of termination domains for contralateral and ipsilateral RGC axons within the dLGN (Huberman et al., 2002; Muir-Robinson et al., 2002). Previous studies showed that expression gradients of ephrin-A proteins across the dLGN, which play a key role in topographic mapping of the nasotemporal axis of the retina onto the dLGN, are also required for proper placement of eye-specific inputs in the dLGN (Huberman et al., 2005; Pfeiffenberger et al., 2005). Teneurin-3, another transmembrane protein expressed in a gradient in the dLGN, was also shown to regulate mapping of ipsilateral retinogeniculate projection (Leamey et al., 2007).

A relatively simple model that could explain the stereotypical placement of the eye-specific domains is that layer-specific positional cues that can discriminate contralateral and ipsilateral RGC axons guide them to the correct eye-specific layers in the dLGN (Chapman, 2000; Crowley and Katz, 1999; Huberman et al., 2002; Shatz, 1996). However, the molecular nature of such layer-specific guidance cues has remained to be elucidated.

Nell2 (<u>N</u>eural <u>e</u>pidermal growth factor (EGF)-l<u>i</u>ke-l<u>i</u>ke 2; initially identified in the chick and named “Nel”) is an extracellular glycoprotein that has structural similarities with thrombospondin-1 and is predominantly expressed in the nervous system (Kuroda et al., 1999; Matsuhashi et al., 1995; Matsuhashi et al., 1996; Nelson et al., 2004; Nelson et al., 2002; Oyasu et al., 2000; Watanabe et al., 1996). We have previously demonstrated that Nell2 acts as an inhibitory guidance cue for chick RGC axons *in vitro* (Jiang et al., 2009; Nakamura et al., 2012).

In this study, we examined functions of Nell2 in the eye-specific retinogeniculate projection *in vivo* and in regulation of ipsilateral and contralateral RGC axon behaviour *in vitro*. Here we show that Nell2 is expressed in the dorsomedial region of the dLGN, which corresponds to the layer receiving ipsilateral RGC axons. *In vivo* axon tracing analyses showed that the eye-specific pattern of the retinogeniculate projection was disrupted in Nell2 null (Nell2^-/-^) mice: Contralateral RGC axons abnormally invaded the ipsilateral layer of the dLGN, whereas ipsilateral axons terminated in partially fragmented patches, thus forming a mosaic pattern of contralateral and ipsilateral axon termination zones. *In vitro,* Nell2 inhibited outgrowth, induced growth cone collapse, and caused repulsion in contralateral, but not ipsilateral, RGC axons. These results provide evidence that Nell2 acts as an inhibitory guidance molecule specific for contralateral RGC axons and prevents them from invading the ipsilateral layer of the dLGN. In addition, our findings indicate that a layer-specific positional label that exerts discriminatory effects on contralateral and ipsilateral axons plays essential roles in establishment of the eye-specific patterns in the visual system.

## RESULTS

### Nell2 expression domain overlaps with the ipsilateral layer of the dLGN

To explore potential functions of Nell2 in development of the eye-specific targeting patterns, we first examined Nell2 expression in the dLGN at the time of retinogeniculate projection by RNA *in situ* hybridization. In the mouse, first contralateral RGC axons enter the dLGN by embryonic day (E) 16, and ipsilateral axons around E18-birth (Godement et al., 1984). Nell2 expression was detected in the developing dLGN at E18.5 (**Fig. 1A**). Between postnatal day 1 (P1) and P10, by when the initial pattern of the retinogeniculate projection is established (Godement et al., 1984; Huberman et al., 2010), Nell2 expression levels increased and became restricted in the dorsomedial region of the dLGN, where ipsilateral RGC axons normally terminate (“ipsilateral layer”) (**Fig. 1B-E**). No obvious gradient of expression was observed for Nell2 in the dLGN (Flanagan, 2006).

**Fig. 1.**
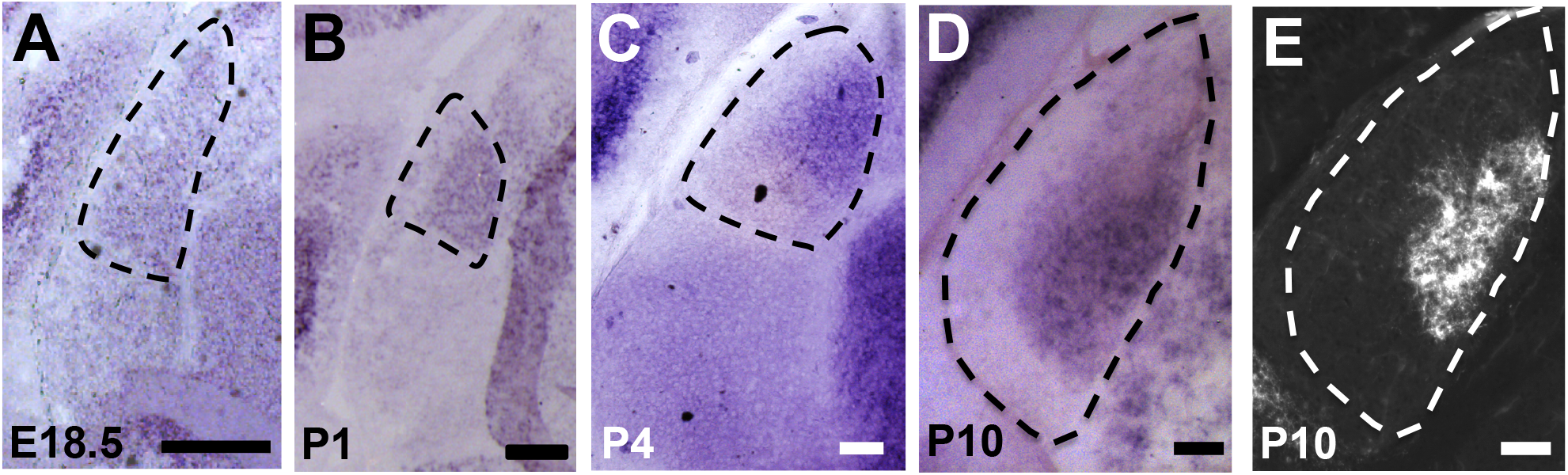
Nell2 expression in the developing mouse dLGN. Coronal sections through the dLGN prepared from E18.5 (A), P1 (B), P4 (C), and P10 (D, E) mouse embryos are shown. Dorsal is at the top, and lateral is on the left. Dotted lines indicate the boundary of dLGN. (A-D) Sections were subjected to RNA *in situ* hybridisation using a probe for Nell2. (E) Termination area of ipsilateral RGC axons in the dLGN at P10. RGC axons were labeled with cholera-toxin B (CTB)-Alexa Fluor at P7. Strong expression of Nell2 was detected in the dorsomedial region of the dLGN, which overlaps with the ipsilateral RGC axon termination zone. Scale bars, 100 μm.

### Nell null mice have defects in the eye-specific retinogeniculate projection

To determine whether Nell2 is involved in development of the eye-specific projection, we examined the retinogeniculate projection in Nell2^-/-^ mice (Matsuyama et al., 2004; Nakamoto et al., 2014). Overall structure and cytoarchitecture of the visual system (retina, optic chiasm, and dLGN) are maintained in the mutant mice (**Figs. 2A, 3, 4**). We labelled the whole eyes with an axon tracer dye of two different colours, and the location of each eye’s projection was examined in the dLGN (Bjartmar et al., 2006; Huberman et al., 2002). As previously reported, in wild-type mice at P12, contralateral and ipsilateral RGC axons terminated in two non-overlapping layers: Ipsilateral axons are confined to a small single patch located in the dorsomedial part of the dLGN, whereas contralateral axons terminated in the surrounding areas (Rossi et al., 2001) (**Fig. 2A**). In contrast, in Nell2^-/-^ mice contralateral axons invaded the ipsilateral layer of the dLGN, and ipsilateral axons terminated in partially fragmented patches, forming a mosaic pattern of contralateral and ipsilateral axon termination zones (**Fig. 2B-D, Supplementary Fig. 1**). Aberrant invasion of the ipsilateral layer by contralateral axons was observed in 10/12 (83.3%) of Nell2^-/-^ mice, but not in wild-type (0/10) mice (p<0.001). Although contralateral and ipsilateral axons terminated in aberrant locations of the dLGN, the segregation of inputs from the right and left eyes occurred properly (**Fig. 2E and Supplementary Fig. 2**). These results indicate that Nell2 is essential for eye-specific targeting of RGC axons but not for segregation of retinal inputs in the dLGN.

**Fig. 2.**
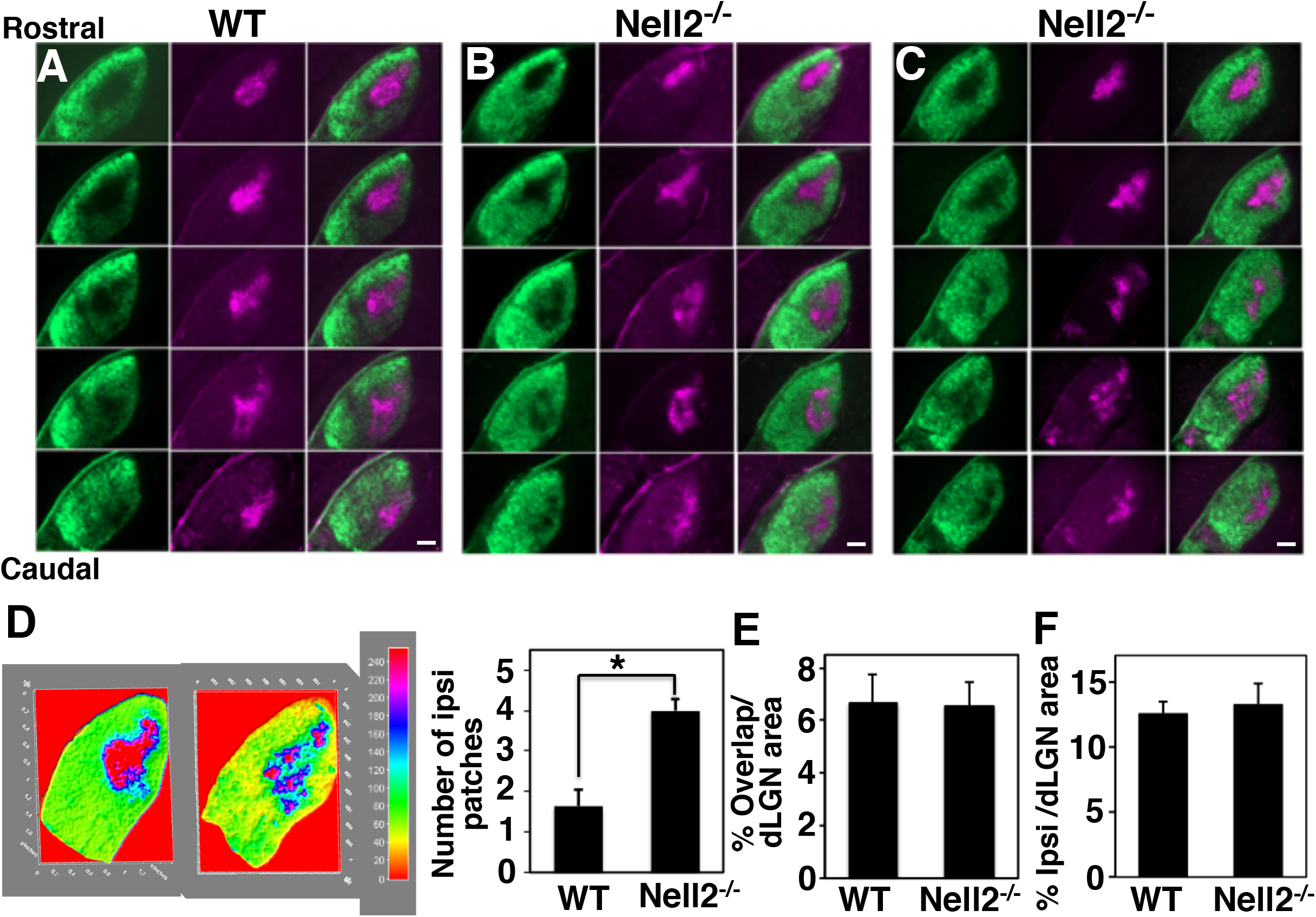
Defects in the eye-specific retinogeniculate projection in *Nell2^-/-^* mice. RGC axons of the right and left eyes were labeled at P9 by injection with cholera toxin B (CTB)-Alexa Fluor 488 (green) and CTB-Alexa Fluor 594 (magenta), respectively. Serial coronal sections through the left dLGN prepared at P12 are shown. In each panel, dorsal is at the top and lateral is on the left. (A) In the wild type (n=10), ipsilateral axons (magenta) are confined to a single patch in a dorsomedial region of the dLGN, and contralateral axons (green) terminate in the surrounding areas. (B and C) Two representative examples of the retinogeniculate projection in Nell2”“ mice (n=12). Contralateral axons invade the ipsilateral layer, forming a mosaic pattern of contralateral and ipsilateral axons (10/12). Scale bars, 100 μm. (d) Quantification of the number of ipsilateral patches. The ipsilateral projection in Nell2^-^ mice appeared patchy compared to that in wild type mice. (e) Contralateral and ipsilateral axon overlap was quantified and presented as the percentage of the total dLGN area containing overlap. (f) Quantification of the ratio of percent ipsilateral projection area to total dLGN area. In (d-f), data are plotted as mean ± s.e.m. *, p<0.0001.

### Nell null mice show no obvious defects in RGC axon pathfinding

It is possible that the aberrant termination patterns observed in Nell2^-/-^ dLGN were caused by misrouting of RGC axons at the optic chiasm. Axons that normally remain ipsilateral might instead cross the midline and project to their normal ‘ipsilateral’ layer in the dLGN, resulting in invasion of the ipsilateral layer by ‘contralateral’ axons. If this were the case, there would be a decreased number of ipsilateral axons in the dLGN. We therefore compared the ratio of ipsilateral patch area to total dLGN area. No significant difference was observed in the size of ipsilateral patch between the wild type and Nell2^-/-^ mice (**Fig. 2F and Supplementary Fig. 3**). We also prepared sections through the optic chiasm region and examined patterns of RGC axon routing in Nell2^-/-^ mice. No obvious defects were observed in the decussation pattern or structure of the optic chiasm, or in RGC axon fasciculation (**Fig. 3**).

**Fig. 3.**
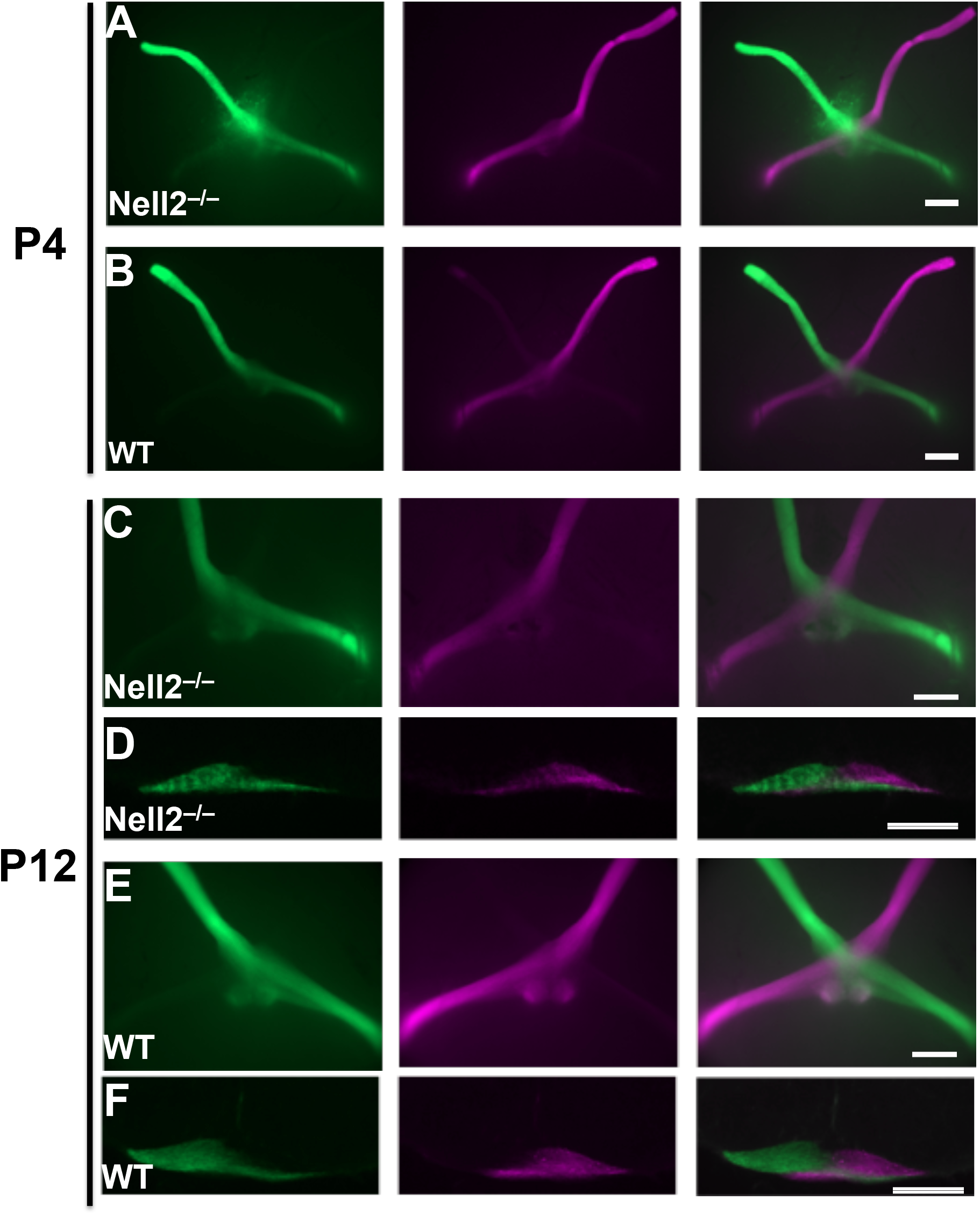
No obvious defects in the decussation patterns at the optic chiasm of *Nell2^-/-^* mice. RGC axons of the right and left eyes were labeled by injection with cholera toxin B (CTB)-Alexa Fluor 488 (green) and CTB-Alexa Fluor 594 (magenta), respectively. The brain was dissected out three day later ((A and B) P4. (C-E) P12), and decussation patterns of the optic nerve at the optic chiasm were observed in whole-mount preparations (A-C, E; ventral views, rostral is at the top) and in coronal sections (d and f; dorsal is at the top). (A, C, D) Nell2^-/-^. (B, E, F) Wild type. Scale bars, 100 μm.

It has been previously reported that Nell2 is expressed in developing RGCs (Nakamoto et al., 2014; Nelson et al., 2002; Wang et al., 2007). It is possible that Nell2 is expressed specifically in contra- or ipsi-laterally projecting RGCs and is involved in specification of their laterality of projection. As shown in **Fig. 4**, however, expression of Nell2 in RGCs is detected throughout the retina, and its expression domain extends to the retinal periphery, including the ventrotemporal region that contains ipsilaterally-projecting RGCs. No obvious gradient of Nell2 expression was observed in the retina. The expression pattern of Nell2 in the retina therefore does not appear to correlate with a particular retinogeniculate projection phenotype. We also found no significant differences in expression patterns of the Zic2 gene, the key transcriptional regulator for determination of ipsilateral RGCs, between wild-type and Nell2^-/-^ mice. Taken together, these results suggest that Nell2^-/-^ mice have no obvious defects in RGC axon pathfinding.

**Fig. 4.**
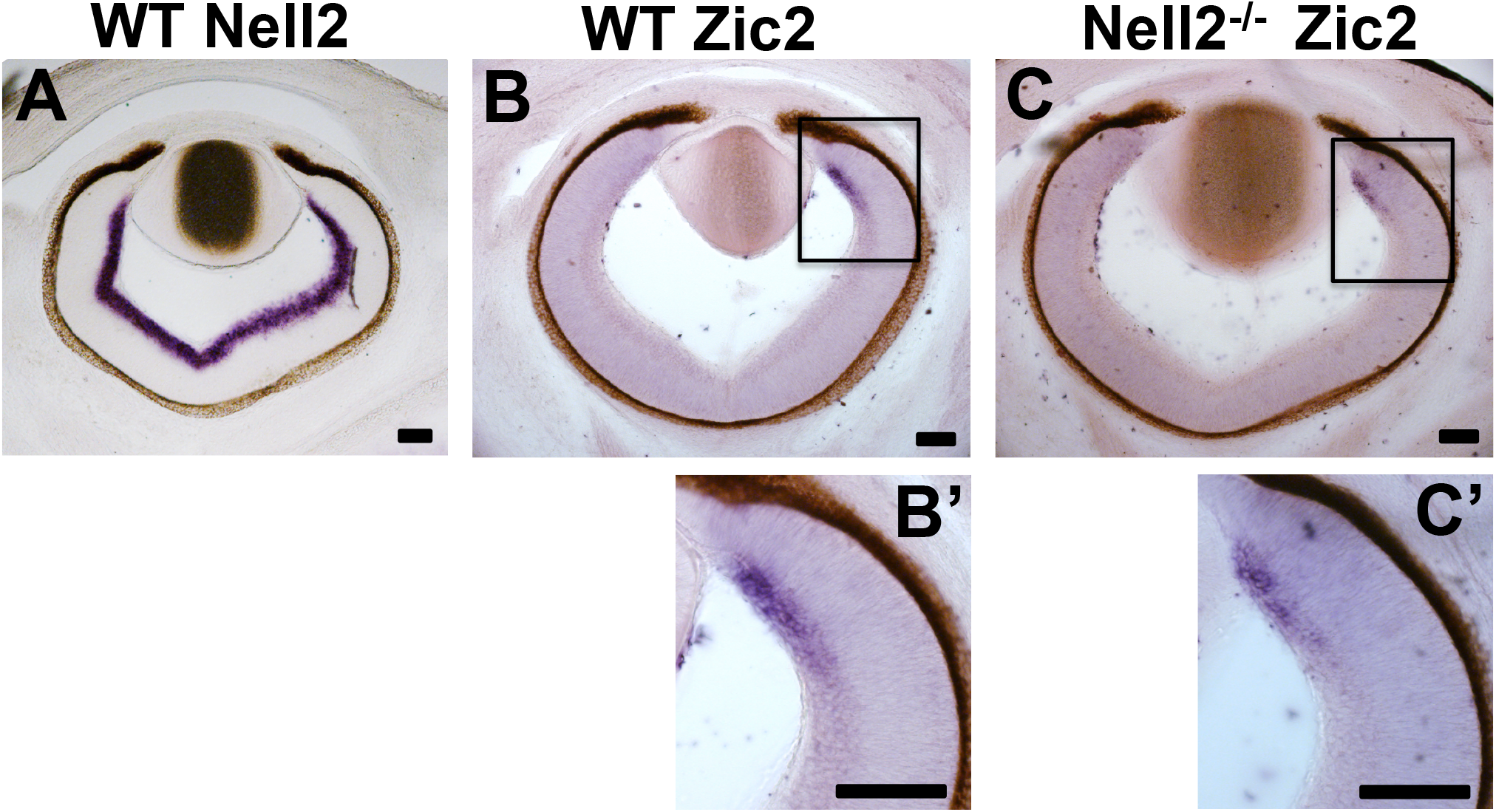
Nell2 and Zic2 expression in the developing mouse retina. Horizontal sections across the ventral retina were prepared from wild type (A, B) or Nell2^-/-^ (C) mouse embryos at E15.5 and hybridised with an RNA probe for Nell2 (A) or Zic2 (B, C). A higher magnification view of the area outlined by the black rectangle in (B) and (C) are shown in (B’) and (C’), respectively. In the wild type, Nell2 is expressed in RGCs throughout the retina, and its expression domain extends to the periphery of the retina (A), including the Zic2-positive domain (B). Zic2 expression is normally confined to the ventrotemporal part of the retina, which contains ipsilaterally-projecting RGCs. No significant alteration in Zic2 expression was observed in Nell2^-/-^ retinae. Scale bars, 200 μm.

### Nell2 inhibits contralateral but not ipsilateral RGC axons *in vitro*

The Nell2 expression domain in the dLGN overlaps with the termination zone of ipsilateral RGC axons, which contralateral axons normally avoid (**Fig. 1**). In addition, we previously demonstrated that Nell2 (Nel) inhibits outgrowth of RGC axons and induces growth cone collapse and axon retraction in the chick (Jiang et al., 2009; Nakamura et al., 2012), in which all the mature RGCs send their projections contralaterally (Thanos and Bonhoeffer, 1984). These findings may suggest that Nell2 acts as an inhibitory guidance cue for contralateral RGC axons and prevents them from terminating in the ipsilateral layer of the dLGN. We therefore examined the effects of Nell2 on contralateral and ipsilateral RGC axons by using several axon behaviour assays *in vitro*.

We first tested effects of Nell2 on RGC axon outgrowth, using retinal explants prepared from the ventrotemporal (VT, containing ipsilaterally projecting RGCs) and ventronasal (VN, contralaterally projecting RGCs) retinae. VT and VN explants were individually cultured on the substratum coated with either a Nell2 protein conjugated with an alkaline phosphatase (AP) tag (Nell2-AP) or control unconjugated AP. We found that Nell2 specifically inhibited outgrowth of RGC axons from VN explants (**Fig. 5**). Modest inhibition of axon outgrowth was also observed for VT explants treated with Nell2-AP, but it was not statistically significant. These results indicate that Nell2 inhibits outgrowth of contralateral RGC axons.

**Fig. 5.**
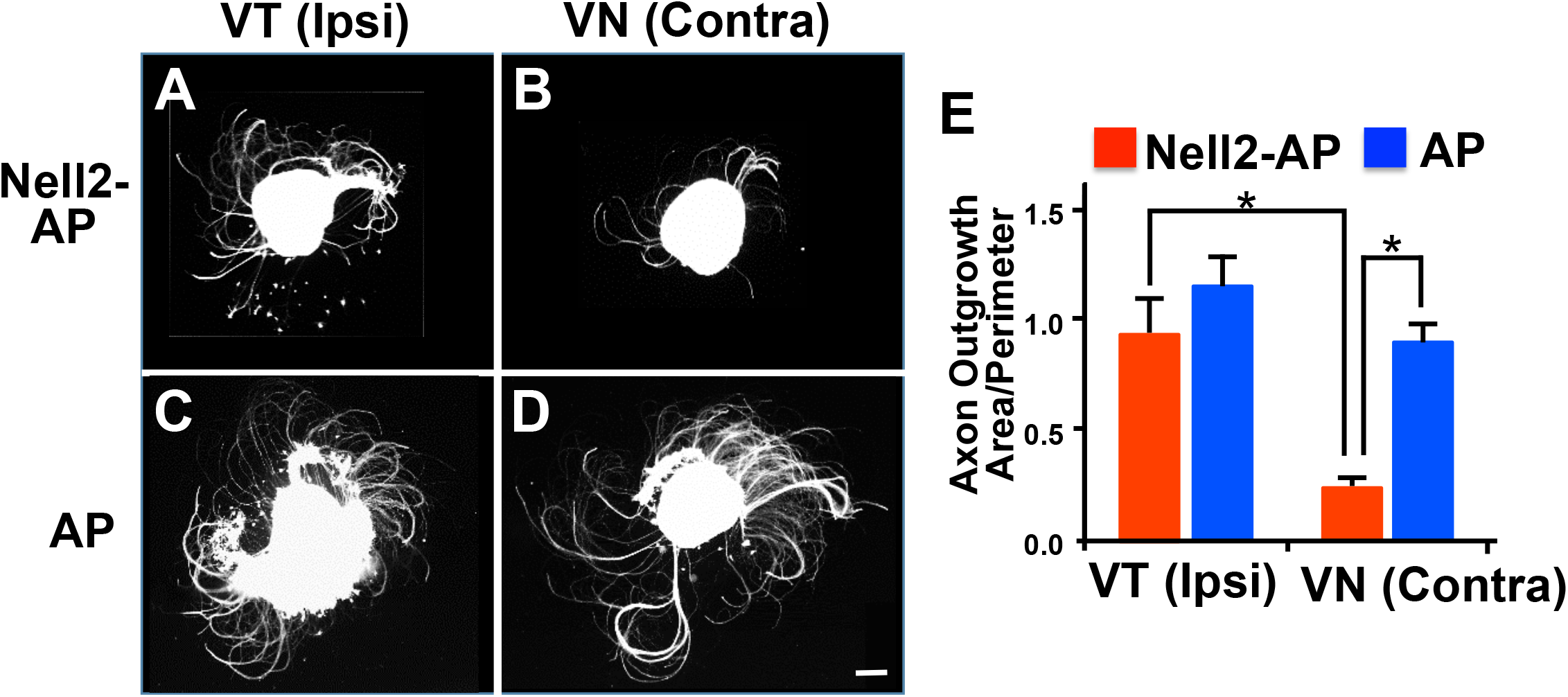
Nell2 inhibits outgrowth of contralaterally projecting RGC axons. Explants of the ventrotemporal (VT; containing ipsilaterally projecting RGCs) (A, C) and ventronasal (VN; contralaterally projecting RGCs) (B, D) retina were prepared from E15.5 mouse embryos and cultured for 72 hours on a substratum coated with Nell2-AP (A, B) or AP (C, D), and axon outgrowth was quantified (E) (n=4 for each condition). Nell2-AP significantly inhibited outgrowth of VN retinal axons. Scale bar, 100 μm. *, p<0.01.

The ability to induce growth cone collapse is a hallmark of many inhibitory axon guidance cues, such as ephrins, slits, and semaphorins. Therefore, next we examined how Nell2 regulates growth cone morphology of RGC axons. We cultured VT and VN retinal explants on a permissive substratum to allow formation of well-developed growth cones at the tip of RGC axons. Then the axons were treated with Nell2-AP or AP, and their effects on morphology of growth cones were observed. Treatment with control AP did not affect the growth cone morphology in VT or VN axons. In contrast, Nell2-AP induced growth cones collapse in most of VN, but not VT, axons (**Fig. 6**). Our results indicate that Nell2 induces growth cone collapse specifically in contralaterally projecting RGC axons.

**Fig. 6.**
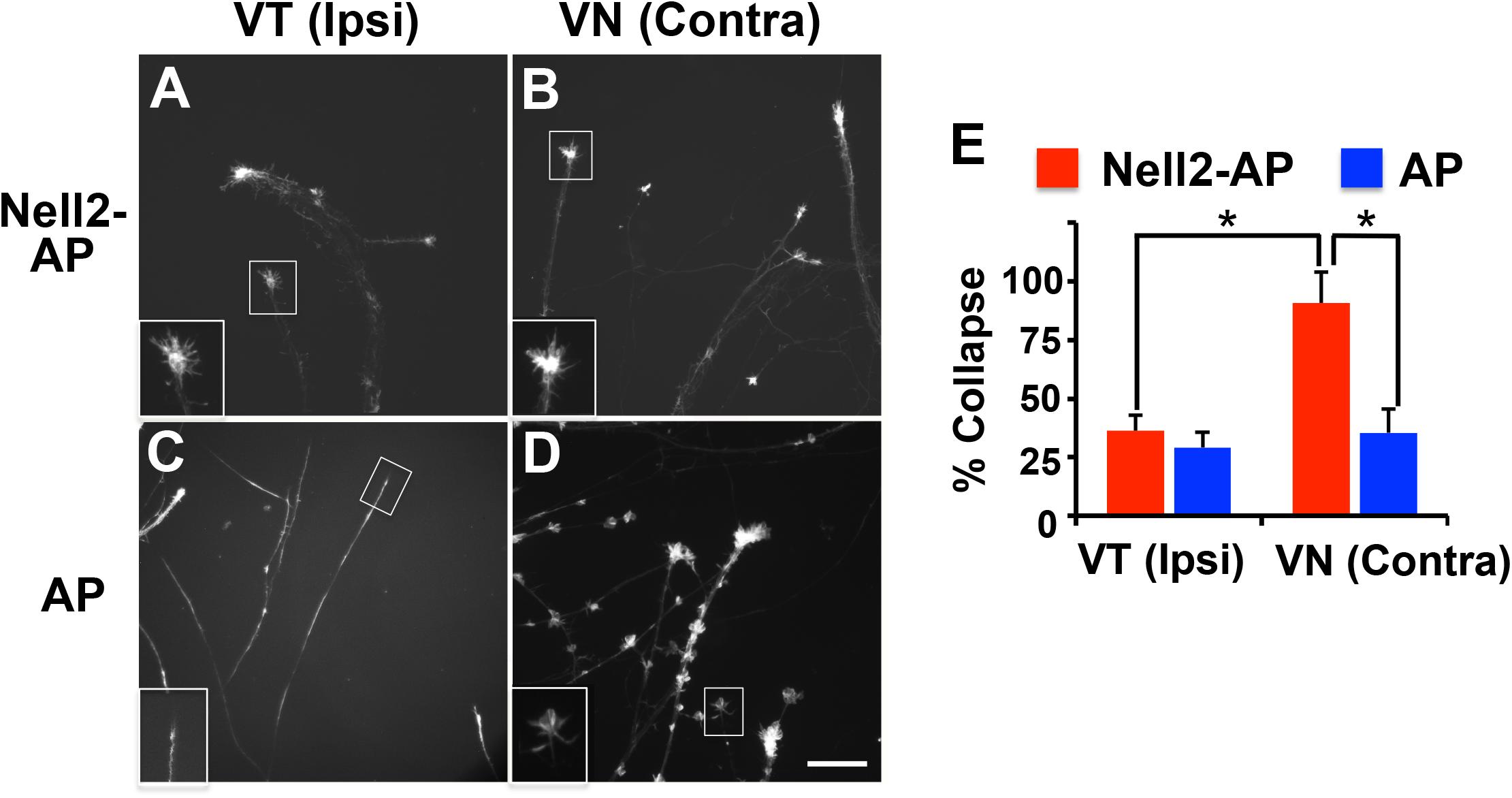
Nell2 induces growth cone collapse specifically in contralaterally projecting RGC axons. VT (A, C) and VN (B, D) retinal explants prepared from E15.5 mouse embryos were cultured *in vitro* for 72 hours and then treated with Nell2-AP (A, B) or AP (C, D) (n=4 for each condition, 30< growth cones were observed in each experiment). The growth cone morphology was observed 30 min later. Higher magnification views of representative growth cone morphology (white squares) are shown in insets. (E) Quantification of the growth cone collapsing activity. The percentage of collapsed growth cones was plotted as mean ± s.e.m. Nell2-AP induced growth cone collapse in VN but not VT retinal axons. Scale bar, 100 μm. *, p<0.001.

We then performed stripe assay (Hornberger et al., 1999) to examine whether contralateral and ipsilateral RGC axons show preference for or against Nell2. VT and VN explants were cultured on the substratum consisting of alternative stripes of Nell2-AP and AP, and thus RGC axons were given a choice to grow on Nell2-AP– or AP–containing substratum. As shown in **Fig. 7**, axons from VN axons prefer to grow on control AP stripes, whereas VT axons did not show preferences and grew randomly. Taken together, these *in vitro* experiments demonstrated that Nell2 acts as a contralateral RGC axon-specific inhibitory guidance cue.

**Fig. 7.**
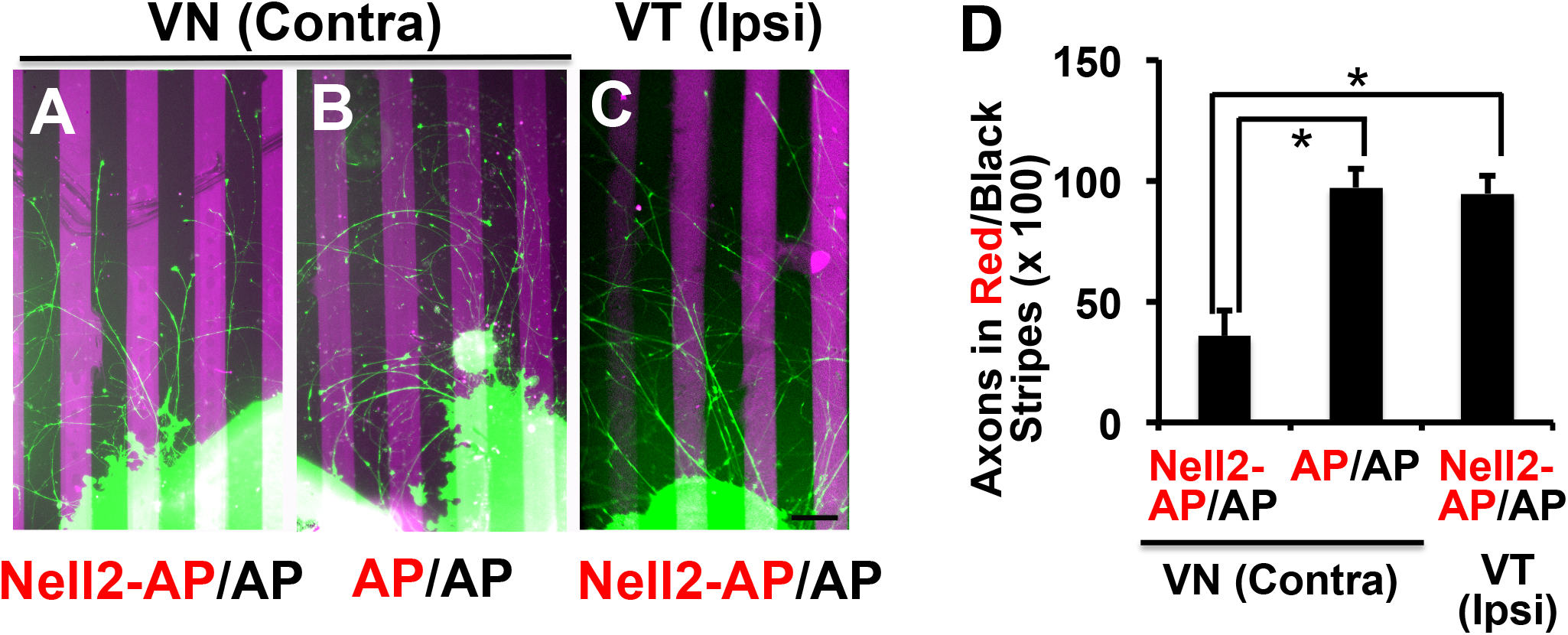
Specific repulsion of contralaterally projecting RGC axons by Nell2. VN (A, B) and VT (C) retinal axons were grown on alternating stripes consisting of Nell2-AP (A, C) or AP (B) containing substrata (labelled with fluorescent microspheres (red)) and AP– containing substrata (black). Patterns of axon outgrowth were observed 72 hours later. VN axons show a strong preference for control AP stripes, whereas VT axons show no preference. (D) Quantification of axon behaviour on Nell2 stripes. Plotted as mean ± s.e.m. n=6 for each experimental condition. Scale bar, 100 μm. *, p<0.001.

## DISCUSSION

It has long been postulated that the eye-specific visual projection is regulated by layer-specific guidance cues; however, their molecular nature has remained elusive. The current study identified Nell2 as a positional label for the ipsilateral layer in the dLGN. In a series of *in vitro* axon behavior assays, Nell2 exerted inhibitory effects specifically on contralateral RGC axons. In Nell2^-/-^ mice, contralateral RGC axons aberrantly invaded the ipsilateral layer of the dLGN. The effects of Nell2 on contralateral RGC axons in the dLGN or *in vitro* are consistent with the model in which Nell2 in the ipsilateral layer acts as an inhibitory guidance cue specific for contralateral RGC axons and prevents them from entering the ipsilateral layer. The removal of the repellent Nell2 allows contralateral axons to invade and arborise ectopically in the ipsilateral layer (**Fig. 8**). Those results are also in agreement with our previous observation in the chick that Nell2 (Nel) inhibits RGC axons (which are all contralaterally-projecting) and is expressed in specific tectal layers that (contralateral) RGC axons do not normally invade (Jiang et al., 2009; Nakamura et al., 2012).

**Fig. 8.**
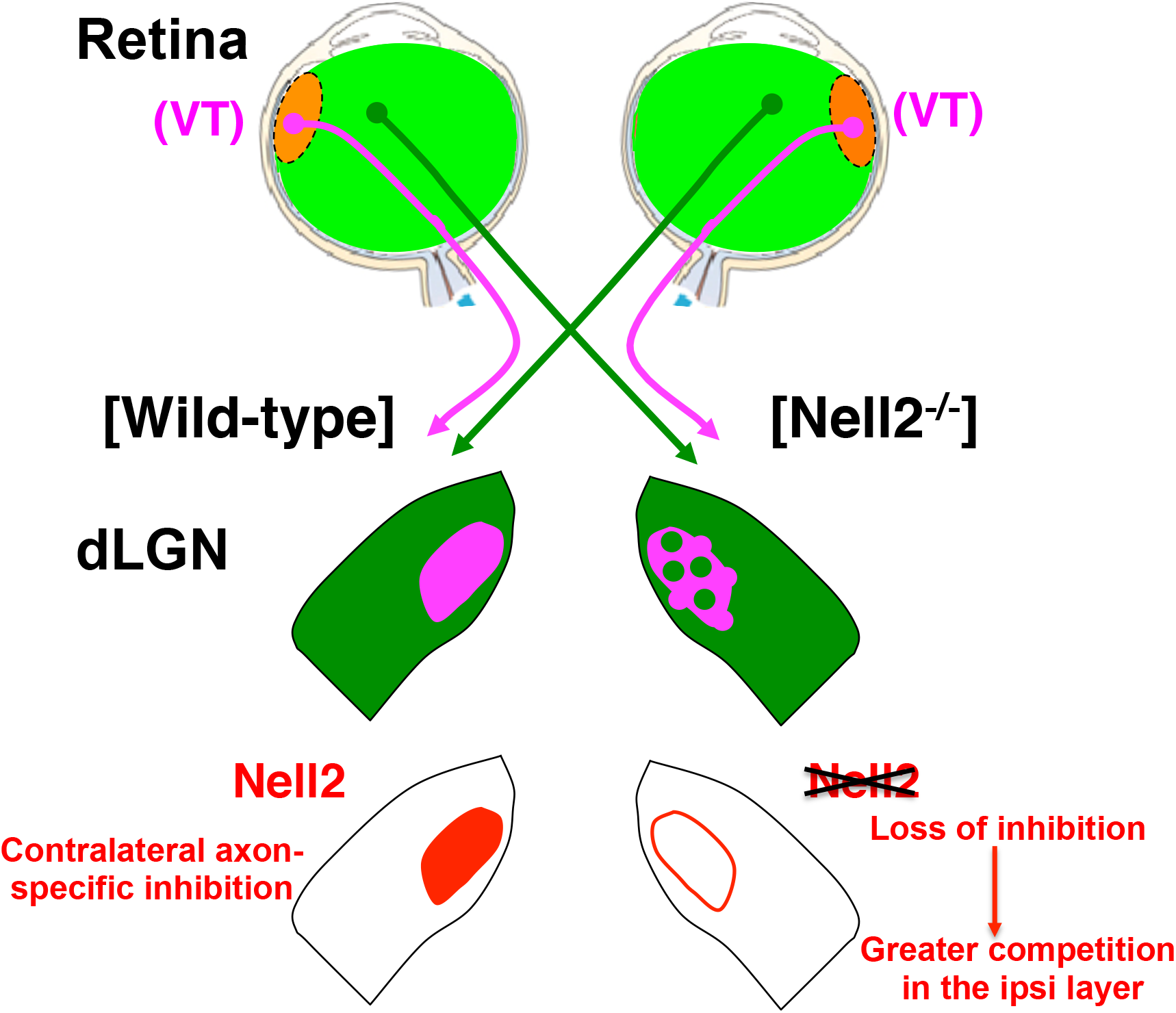
Nell2 and eye-specific retinogeniculate projection. (A) In the mouse visual system, the ipsilateral RGC axons (c.3-5% of the optic nerve fibres) arise from a ventrotemporal (VT) segment of the retina. In the dLGN, ipsilateral axons are confined to a small single patch located in the dorsomedial part of the dLGN (magenta), whereas contralateral axons terminated in the surrounding areas (green). In Nell2^-/-^ mice, contralateral RGC axons abnormally invade the ipsilateral layer, whereas ipsilateral axons terminated in partially fragmented patches, thus forming a mosaic pattern of contralateral and ipsilateral axon termination zones. (B) Repulsion/competition model. Axons compete with one another for space in the target. The contralateral axon-specific repellent Nell2 (red) in the ipsilateral layer biases this competition: Contralateral RGC axons are repelled by Nell2, and thus they are forced to terminate in the surrounding contralateral layer; whereas ipsilateral axons can project to the ipsilateral layer. Ipsilateral axons do not terminate in the contralateral layer because there is greater axon-axon competition, so they prefer to avoid this competition and arborize only in the ipsilateral layer. In Nell2^-/-^ mice, the Nell2 repellent is removed, and contralateral axons can now invade the ipsilateral layer and compete more effectively with ipsilateral axons there. Subsets of the ipsilateral axons lose competition and spread out into surrounding areas in the contralateral layer.

Whereas the expression patterns of Nell2 in the developing mouse dLGN do not show obvious gradients, positional labels expressed in gradients across the projecting and target areas play key roles in topographic neural map formation (Flanagan, 2006). Ephrin-As are expressed in gradients in the dLGN (high in ventral-lateral-anterior and low in dorsal-medial-posterior) and, through interactions with EphA receptors expressed in complementary gradients in the retina (high in temporal, low in nasal), regulate formation of topographic mapping along the nasotemporal axis in the retina onto the dLGN (Drescher et al., 1995; Feldheim et al., 2004; Feldheim et al., 1998; Nakamoto et al., 1996). Interestingly, ephrin-As have also been implicated in eye-specific axon targeting in the dLGN of the mouse (Pfeiffenberger et al., 2005) and ferret (Huberman et al., 2005). Similarly, the cell surface adhesion molecule teneurin-3 is expressed in corresponding gradients in the dLGN (high in dorsal, low in ventral) and the retina (high in ventral, low in dorsal), and regulates the eye-specific patterning of the retinogeniculate projection through homophilic interactions (Leamey et al., 2007). Considering that the ipsilaterally-projecting RGCs localise in the ventrotemporal region in the retina, it seems plausible that the key contributing factors in ephrin-A– and teneurin-3–mediated patterning of eye-specific retinogeniculate projections are the nasal-retina versus temporal-retina distinction and the ventral-retina versus dorsal-retina distinction, respectively (Huberman et al., 2005). In contrast, our results in this study suggest that Nell2 contributes to establish the eye-specific projection patterns by utilizing the contralateral-retina versus ipsilateral-retina distinction.

It is noteworthy that deletion of Nell2 also affected ipsilateral RGC axons in the dLGN. In Nell2^-/-^ mice, subsets of ipsilateral axons appeared to be displaced from the ipsilateral layer and to terminate in partially fragmented patches in the dLGN. This phenotype could not be simply explained by inhibitory effects of Nell2 on contralateral axons alone. One model that could account for the behaviour of ipsilateral axons in the Nell2^-/-^ dLGN involves axon-axon competition, in combination with contralateral axon-specific repulsion by Nell2. It is thought that during development axons compete with one another to fill available space in the target region (Holt and Harris, 1993; Jacobson and Rao, 2005). In normal development, by acting as a repellent for contralateral RGC axons, Nell2 would bias this competition and give ipsilateral axons an advantage in the Nell2-containing ipsilateral layer: Contralateral RGC axons are repelled by Nell2 in the ipsilateral layer, and thus they would be forced to terminate in the surrounding contralateral layer; whereas ipsilateral axons are not repelled by Nell2 and can terminate in the ipsilateral layer. Ipsilateral axons do not terminate in the contralateral layer because there is greater axon-axon competition, so they prefer to avoid this competition and arborise only in the ipsilateral layer. In Nell2^-/-^ mice, the Nell2 repellent is removed, and thus contralateral axons are now able to invade the ipsilateral layer and compete more effectively with ipsilateral axons there. Ipsilateral axons face increased competition in the ipsilateral layer, and subsets of the ipsilateral axons lose competition and spread out into surrounding areas in the contralateral layer. The model thus seems to fit well with the phenotypes of contra- and ipsi-lateral RGC axons in Nell2^-/-^ mice observed in this study. Similar repulsion/competition models in which axon-axon competition is biased by non-uniform expression of guidance molecules in the target region were proposed to explain the aberrant projection patterns of nasal RGC axons in ephrin-A mutant mice (Feldheim et al., 2000; Feldheim et al., 1998).

Despite the aberrant patterns of contra- and ipsi-lateral axon termination, inputs from the right and left eye are still segregated, suggesting that Nell2 is required for proper placement of eye-specific inputs in the dLGN but is not essential for segregation of those inputs. Similar results have been previously reported for ephrin-As: Gradients of ephrin-As in the dLGN are required for the proper placement of contra- and ipsi-lateral inputs but not 0required for their segregations (Pfeiffenberger et al., 2005). Taken together, those findings indicate differential and complementary contributions of activity-dependent and activity-independent mechanisms to the establishment of the eye-specific retinogeniculate projection.

Recently Nell2 has been shown to repel murine spinal commissural axons through the Robo3 receptor and steer them towards and across the midline of the spinal cord (Jaworski et al., 2015). In the developing mouse retina, however, Robo3 is not expressed in RGCs (Blackshaw et al., 2004); and expression of related Robo2 and Robo1 are detected throughout the RGC layer (Robo2 is expressed in most cells and Robo1 in a scattered subpopulation of cells in the RGC layer) and does not correlate with a particular retinogeniculate projection phenotype (Erskine et al., 2000). Therefore it seems unlikely that Robo receptors are responsible for the Nell2-medicated eye-specific retinogeniculate projection. In addition, whereas EGF-like repeats of the Nell2 protein appear to be responsible for repulsion of spinal commissural axons (Jaworski et al., 2015), our previous studies have revealed that cysteine-rich domains exert inhibition of retinal axons (Nakamoto et al., 2014). These findings suggest that Nell2 regulates behaviour of retinal and spinal commissural axons through its different domains binding to distinct cognate receptors. Interestingly, it was shown that different receptors mediate VEGF-A–induced attraction in RGC axons (the NRP1 receptor) and spinal commissural axons (the FLK1 (KDR/VEGFR-2) receptor) (Erskine et al., 2011; Ruiz de Almodovar et al., 2011). In view of the structural similarities between Nell2 and thrombospondin-1, it seems likely that Nell2 interacts with a diverse range of cell surface molecules by using different domains. Identification of functional Nell2 receptors for RGCs axon guidance will be required to fully understand signalling mechanisms for Nell2-mediated eye-specific visual projection.

## MATERIALS AND METHODS

### Nell2 mutant mice and genotyping

Mutant mice carrying the Nell2 null allele have been described previously (Matsuyama et al., 2004) and were maintained on a C57BL/6 background in the Medical Research Facility at the University of Aberdeen. The animals were used under licenses from the UK Home Office in accordance with the Animals (Scientific Procedures) Act 1986 and following approval from the University’s Ethical Review Committee. For genotyping, genomic DNA was PCR-amplified using primers (5’-ATGGAATCCCGGGTGTTACT-3’) and (5’-CTCCCCAAGTTCTAACTCTG-3’) for the wild-type Nell2 locus, and (5’-ATGTATGGTGGTGGGAGGATGC-3’) and (5’-GCCTTCTTGACGAGTTCTTCTGA-3’) for the mutant allele.

### RNA *in situ* hybridization and immunohistochemistry

*In situ* RNA hybridization on mouse brain sections (30-50 μm) was performed by using digoxigenin-labeled probes for Nell2 as previously described (Nakamura et al., 2012).

### Retinal axon tracing

Mouse pups were anesthetized by isoflurane inhalation and received intravitreal injections of cholera toxin-β subunit (CTβ) conjugated to Alexa 488 dye (green label) into one eye and CTβ conjugated to Alexa 594 dye (red label) into the other eye (2–3 μl per eye; 0.5% in sterile saline; Invitrogen, Eugene, OR). Three days later, brains were dissected out, fixed with 4% paraformaldehyde overnight, and embedded in 3% low melting point agarose (Sigma, St. Louis, MO) in PBS. Then coronal sections (50 μm) were cut using a vibratome (Leica).

Images of retinal axon projections from the two eyes were captured with a CCD camera attached to a Zeiss (Thornwood, NY) with 10× and 20× objectives and digitized independently using Image J. For quantification of ipsilateral patches, only three sections that contained the largest ipsilateral projections (corresponding the middle third of the LGN) were analysed. For quantification of eye-specific segregation, the boundary of the dLGN was outlined, excluding the intrageniculate leaflet, the ventral LGN, and the optic tract. The pixel overlap between ipsilateral and contralateral projections was measured, and the proportion of dLGN occupied by ipsilateral axons was measured as a ratio of ipsilateral pixels to the total number of pixels in the dLGN region.

### Axon outgrowth assays

Retinal explants were prepared from VT and VN areas of the E15.5 mouse retinae. The explants were cultured in a 4-well culture dish (Nunc) pre-coated with 50 μg/ml of laminin and then with 0.25 μM of Nell2-AP or AP. Explants were cultured in the retinal culture medium (15% fetal bovine serum, 0.6% glucose, penicillin/streptomycin in DMEM:F 12=1:1). After 48-72 hours, axons were labelled by incubating the cultures in 33 μM carboxylfluorescein diacetate succinimidyl ester (Molecular Probes, Eugene, OR) for 5 minutes and photographed. For quantification of axon outgrowth, areas of axon growth were measured and normalized by the length of their perimeter using Image J. Axon outgrowth was expressed as the mean ± s.e.m.

### Growth Cone Collapse Assays

Growth cone collapse assays were performed essentially as described previously (Nakamura et al., 2012). Retinal explants were prepared from E15.5 mouse embryos and cultured for 48-72 hours on the substratum coated with 100 μg/ml laminin and in the retinal culture medium. Then retinal axons were treated with 0.25 μM Nell2-AP or control AP. The explants were incubated at 37°C for up to 30 min, fixed, and stained with Alexa Fluor 488 Phalloidin (Invitrogen). Individual growth cones were scored as collapsed or not collapsed, and percentages of collapsed growth cones were calculated. In each experiment, at least 30 growth cones for each condition were scored, and three independent experiments were performed.

### Stripe Assays

Stripe assays using purified Nell2-AP and unconjugated AP protein were performed as previously described (Knoll et al., 2007). Briefly, a silicon matrix with 90 μm channels (obtained from the laboratory of Prof. Martin Bastmeyer (Zoologisches Institut, Universität Karlsruhe, Germany)) was placed onto a 6 cm petri dish, applied with a first protein solution (0.5 μM Nell2 or AP conjugated with AlexaFluoro 647), and incubated for 30 min at 37°C in a moist chamber to set the first stripes. After rinsing the channels with HBSS, the matrix was removed. A second protein solution (0.5 μM AP) was applied to the stripe area, and the dishes were incubated for 30 min at 37°C. The second protein solution was then removed and the dishes were rinsed with HBSS. Then the stripes were coated with 20 μg/ml laminin in HBSS (Invitrogen Cat 23017015) for 2 hrs at 37°C in a moist chamber, and then rinsed with HBSS. Retinal explants prepared from E15.5 mouse embryos were cultured on the stripes in the retinal culture medium for 2~3 days. Axons were stained with Alexa flour 488 Phalloidin. To generate a quantitative index of RGC axon growth on membrane stripes, the integrated fluorescent signal from the axons in the first stripes versus that in the second stripes was calculated for a rectangular region of interest that spanned the width of the image adjacent to the explant edge by using Image J (Stettler et al., 2012).

### Statistical analysis

For statistical analysis of the retinogeniculate projection patterns in Nell2 mutant mice, an unpaired Student’s t test was used. Results of the *in vitro* axon behaviour assays were analysed by ANOVA. P values are given in the figure legends. No statistical methods were used to predetermine the sample sizes, but our sample sizes are similar to those generally employed in the field. Part of data collection and analysis were performed blind to the conditions of the experiments.

## Acknowledgements

We thank Lynda Erskine, Colin McCaig, and David Feldheim for discussion and comments on the manuscript; the Imaging Core Facility of the Institute of Medical Sciences and the Medical Research Facility of University of Aberdeen for technical supports. This work was supported by grants from Biotechnology and Biological Sciences Research Council (BB/G007632/1) and Royal Society (MN).

## Author contributions

C. N. and M.N. conceived the project and designed the research; C.N., E.D., and M.N. performed the experiments and analysed the data; M.H. provided Nell2 mutant mice and advice on their maintenance. C.N. and M.N. wrote the manuscript with contributions from all authors.

## REFERENCES

Bjartmar, L., Huberman, A. D., Ullian, E. M., Renteria, R. C., Liu, X., Xu, W., Prezioso, J., Susman, M. W., Stellwagen, D., Stokes, C. C., et al. (2006). Neuronal pentraxins mediate synaptic refinement in the developing visual system. J. Neurosci. 26, 6269–6281.

Blackshaw, S., Harpavat, S., Trimarchi, J., Cai, L., Huang, H., Kuo, W. P., Weber, G., Lee, K., Fraioli, R. E., Cho, S. H., et al. (2004). Genomic analysis of mouse retinal development. PLoS Biol 2, E247.

Chapman, B. (2000). Necessity for afferent activity to maintain eye-specific segregation in ferret lateral geniculate nucleus. Science 287, 2479–2482.

Crowley, J. C. and Katz, L. C. (1999). Development of ocular dominance columns in the absence of retinal input. Nat. Neurosci. 2, 1125–1130.

Drescher, U., Kremoser, C., Handwerker, C., Loschinger, J., Noda, M. and Bonhoeffer, F. (1995). In vitro guidance of retinal ganglion cell axons by RAGS, a 25 kDa tectal protein related to ligands for Eph receptor tyrosine kinases. Cell 82, 359–370.

Erskine, L. and Herrera, E. (2014). Connecting the retina to the brain. ASN Neuro 6.

Erskine, L., Reijntjes, S., Pratt, T., Denti, L., Schwarz, Q., Vieira, J. M., Alakakone, B., Shewan, D. and Ruhrberg, C. (2011). VEGF signaling through neuropilin 1 guides commissural axon crossing at the optic chiasm. Neuron 70, 951–965.

Erskine, L., Williams, S. E., Brose, K., Kidd, T., Rachel, R. A., Goodman, C. S., Tessier-Lavigne, M. and Mason, C. A. (2000). Retinal ganglion cell axon guidance in the mouse optic chiasm: expression and function of robos and slits. J. Neurosci. 20, 4975–4982.

Feldheim, D. A., Kim, Y. I., Bergemann, A. D., Frisén, J., Barbacid, M. and Flanagan, J. G. (2000). Genetic analysis of ephrin-A2 and ephrin-A5 shows their requirement in multiple aspects of retinocollicular mapping. Neuron 25, 563–574.

Feldheim, D. A., Nakamoto, M., Osterfield, M., Gale, N. W., DeChiara, T. M., Rohatgi, R., Yancopoulos, G. D. and Flanagan, J. G. (2004). Loss-of-function analysis of EphA receptors in retinotectal mapping. J. Neurosci. 24, 2542–2550.

Feldheim, D. A., Vanderhaeghen, P., Hansen, M. J., Frisén, J., Lu, Q., Barbacid, M. and Flanagan, J. G. (1998). Topographic guidance labels in a sensory projection to the forebrain. Neuron 21, 1303–1313.

Flanagan, J. G. (2006). Neural map specification by gradients. Curr. Opin. Neurobiol. 16, 59–66.

Garcia-Frigola, C. and Herrera, E. (2010). Zic2 regulates the expression of Sert to modulate eye-specific refinement at the visual targets. The EMBO journal 29, 3170–3183.

Godement, P., Salaun, J. and Imbert, M. (1984). Prenatal and postnatal development of retinogeniculate and retinocollicular projections in the mouse. J. Comp. Neurol. 230, 552–575.

Heesy, C. P. (2004). On the relationship between orbit orientation and binocular visual field overlap in mammals. Anat Rec A Discov Mol Cell Evol Biol 281, 1104–1110.

Holt, C. E. and Harris, W. A. (1993). Position, guidance, and mapping in the developing visual system. J. Neurobiol. 24, 1400–1422.

Hornberger, M. R., Dutting, D., Ciossek, T., Yamada, T., Handwerker, C., Lang, S., Weth, F., Huf, J., Wessel, R., Logan, C., et al. (1999). Modulation of EphA receptor function by coexpressed ephrinA ligands on retinal ganglion cell axons. Neuron 22, 731–742.

Huberman, A. D. (2007). Mechanisms of eye-specific visual circuit development. Curr. Opin. Neurobiol. 17, 73–80.

Huberman, A. D., Clandinin, T. R. and Baier, H. (2010). Molecular and cellular mechanisms of lamina-specific axon targeting. Cold Spring Harb Perspect Biol 2, a001743.

Huberman, A. D., Feller, M. B. and Chapman, B. (2008). Mechanisms underlying development of visual maps and receptive fields. Annu. Rev. Neurosci. 31, 479–509.

Huberman, A. D., Murray, K. D., Warland, D. K., Feldheim, D. A. and Chapman, B. (2005). Ephrin-As mediate targeting of eye-specific projections to the lateral geniculate nucleus. Nat. Neurosci. 8, 1013–1021.

Huberman, A. D., Stellwagen, D. and Chapman, B. (2002). Decoupling eye-specific segregation from lamination in the lateral geniculate nucleus. J. Neurosci. 22, 9419–9429.

Huh, G. S., Boulanger, L. M., Du, H., Riquelme, P. A., Brotz, T. M. and Shatz, C. J. (2000). Functional requirement for class I MHC in CNS development and plasticity. Science 290, 2155–2159.

Jacobson, M. and Rao, M. S. (2005). Developmental neurobiology (4th edn). New York: Kluwer Academic/Plenum.

Jaworski, A., Tom, I., Tong, R. K., Gildea, H. K., Koch, A. W., Gonzalez, L. C. and Tessier-Lavigne, M. (2015). Operational redundancy in axon guidance through the multifunctional receptor Robo3 and its ligand NELL2. Science 350, 961–965.

Jiang, Y., Obama, H., Kuan, S. L., Nakamura, R., Nakamoto, C., Ouyang, Z. and Nakamoto, M. (2009). In vitro guidance of retinal axons by a tectal lamina-specific glycoprotein Nel. Mol. Cell. Neurosci. 41, 113–119.

Knoll, B., Weinl, C., Nordheim, A. and Bonhoeffer, F. (2007). Stripe assay to examine axonal guidance and cell migration. Nature protocols 2, 1216–1224.

Kuroda, S., Oyasu, M., Kawakami, M., Kanayama, N., Tanizawa, K., Saito, N., Abe, T., Matsuhashi, S. and Ting, K. (1999). Biochemical characterization and expression analysis of neural thrombospondin-1-like proteins NELL1 and NELL2. Biochem. Biophys. Res. Commun. 265, 79–86.

Leamey, C. A., Merlin, S., Lattouf, P., Sawatari, A., Zhou, X., Demel, N., Glendining, K. A., Oohashi, T., Sur, M. and Fassler, R. (2007). Ten_m3 regulates eye-specific patterning in the mammalian visual pathway and is required for binocular vision. PLoS Biol 5, e241.

Matsuhashi, S., Noji, S., Koyama, E., Myokai, F., Ohuchi, H., Taniguchi, S. and Hori, K. (1995). New gene, nel, encoding a M(r) 93 K protein with EGF-like repeats is strongly expressed in neural tissues of early stage chick embryos. Dev. Dyn. 203, 212–222.

Matsuhashi, S., Noji, S., Koyama, E., Myokai, F., Ohuchi, H., Taniguchi, S. and Hori, K. (1996). New gene, nel, encoding a Mr 91 K protein with EGF-like repeats is strongly expressed in neural tissues of early stage chick embryos. Dev. Dyn. 207, 233–234.

Matsuyama, S., Aihara, K., Nishino, N., Takeda, S., Tanizawa, K., Kuroda, S. and Horie, M. (2004). Enhanced long-term potentiation in vivo in dentate gyrus of NELL2-deficient mice. Neuroreport 15, 417–420.

Muir-Robinson, G., Hwang, B. J. and Feller, M. B. (2002). Retinogeniculate axons undergo eye-specific segregation in the absence of eye-specific layers. J. Neurosci. 22, 5259–5264.

Nakamoto, C., Kuan, S. L., Findlay, A. S., Durward, E., Ouyang, Z., Zakrzewska, E. D., Endo, T. and Nakamoto, M. (2014). Nel positively regulates the genesis of retinal ganglion cells by promoting their differentiation and survival during development. Mol. Biol. Cell 25, 234–244.

Nakamoto, M., Cheng, H. J., Friedman, G. C., McLaughlin, T., Hansen, M. J., Yoon, C. H., O’Leary, D. D. and Flanagan, J. G. (1996). Topographically specific effects of ELF-1 on retinal axon guidance in vitro and retinal axon mapping in vivo. Cell 86, 755–766.

Nakamura, R., Nakamoto, C., Obama, H., Durward, E. and Nakamoto, M. (2012). Structure-Function Analysis of Nel, a Thrombospondin-1-like Glycoprotein Involved in Neural Development and Functions. J. Biol. Chem. 287, 3282–3291.

Nelson, B. R., Claes, K., Todd, V., Chaverra, M. and Lefcort, F. (2004). NELL2 promotes motor and sensory neuron differentiation and stimulates mitogenesis in DRG in vivo. Dev. Biol. 270, 322–335.

Nelson, B. R., Matsuhashi, S. and Lefcort, F. (2002). Restricted neural epidermal growth factor-like like 2 (NELL2) expression during muscle and neuronal differentiation. Mech. Dev. 119 Suppl 1, S11–19.

Newton, I. (1730). Opticks or A Treatise of the Reflections, Refractions, Inflections & Colours of Light (Based on the 4th edition edn). New York: Dover Publications, Inc.

O’Leary, D. M., Gerfen, C. R. and Cowan, W. M. (1983). The development and restriction of the ipsilateral retinofugal projection in the chick. Brain Res. 312, 93–109.

Oyasu, M., Kuroda, S., Nakashita, M., Fujimiya, M., Kikkawa, U. and Saito, N. (2000). Immunocytochemical localization of a neuron-specific thrombospondin-1-like protein, NELL2: light and electron microscopic studies in the rat brain. Brain Res. Mol. Brain Res. 76, 151–160.

Petros, T. J., Rebsam, A. and Mason, C. A. (2008). Retinal axon growth at the optic chiasm: to cross or not to cross. Annu. Rev. Neurosci. 31, 295–315.

Pettigrew, J. D. (1986). The evolution of binocular vision. In Vis. Neurosci. (ed. J. D. Pettigrew, K. J. Sanderson & W. R. Levick), pp. 208–222. Cambridge: Cambridge University Press.

Pfeiffenberger, C., Cutforth, T., Woods, G., Yamada, J., Renteria, R. C., Copenhagen, D. R., Flanagan, J. G. and Feldheim, D. A. (2005). Ephrin-As and neural activity are required for eye-specific patterning during retinogeniculate mapping. Nat. Neurosci. 8, 1022–1027.

Polyak, S. (1957). The Vertebrate Visual System. Chicago: University of Chicago Press.

Rossi, F. M., Pizzorusso, T., Porciatti, V., Marubio, L. M., Maffei, L. and Changeux, J. P. (2001). Requirement of the nicotinic acetylcholine receptor beta 2 subunit for the anatomical and functional development of the visual system. Proc. Natl. Acad. Sci. U. S. A. 98, 6453–6458.

Ruiz de Almodovar, C., Fabre, P. J., Knevels, E., Coulon, C., Segura, I., Haddick, P. C., Aerts, L., Delattin, N., Strasser, G., Oh, W. J., et al. (2011). VEGF mediates commissural axon chemoattraction through its receptor Flk1. Neuron 70, 966–978.

Shatz, C. J. (1996). Emergence of order in visual system development. Proc. Natl. Acad. Sci. U. S. A. 93, 602–608.

Stettler, O., Joshi, R. L., Wizenmann, A., Reingruber, J., Holcman, D., Bouillot, C., Castagner, F., Prochiantz, A. and Moya, K. L. (2012). Engrailed homeoprotein recruits the adenosine A1 receptor to potentiate ephrin A5 function in retinal growth cones. Development 139, 215–224.

Thanos, S. and Bonhoeffer, F. (1984). Development of the transient ipsilateral retinotectal projection in the chick embryo: a numerical fluorescence-microscopic analysis. J. Comp. Neurol. 224, 407–414.

Wang, J. T., Kunzevitzky, N. J., Dugas, J. C., Cameron, M., Barres, B. A. and Goldberg, J. L. (2007). Disease gene candidates revealed by expression profiling of retinal ganglion cell development. J. Neurosci. 27, 8593–8603.

Watanabe, T. K., Katagiri, T., Suzuki, M., Shimizu, F., Fujiwara, T., Kanemoto, N., Nakamura, Y., Hirai, Y., Maekawa, H. and Takahashi, E. (1996). Cloning and characterization of two novel human cDNAs (NELL1 and NELL2) encoding proteins with six EGF-like repeats. Genomics 38, 273–276.

Wilks, T. A., Harvey, A. R. and Rodger, J. (2013). Seeing with Two Eyes: Integration of Binocular Retinal Projections in the Brain. In Functional Brain Mapping and the Endeavor to Understand the Working Brain (ed. F. Signorelli & D. Chirchiglia), pp. 227–250. Rijecka, Croatia: InTech.

